# Blood-brain barrier permeability to enrofloxacin in Pengze crucian carp (*Carassius auratus var*. Pengze)

**DOI:** 10.1101/2022.02.27.482137

**Authors:** Fan Zhang, Runping Wang, Jianzhen Huang, Haixin Zhang, Long Wang, Zhiwei Zhong, Jiming Ruan, Huazhong Liu

**Author notes:** Correspondence: Jiming Ruan, Department of Aquaculture, College of Animal Science & Technology, Jiangxi Agricultural University, Nanchang 330045, P.R. China. Huazhong Liu, College of Chemistry & Environmental Science, Guangdong Ocean University, Zhanjiang 524088, P.R. China.

## Abstract

To explore the permeability of blood-brain barrier (BBB) to enrofloxacin (ENR) and the role of brain injury markers in the evaluation of brain damage, crucian carp (*Carassius auratus var*. Pengze) was employed in this work as the research object treated by a single oral administration of the antibiotic. Results showed that ENR residues could be detected in brain of crucian carp exposed to both half lethal dose (LD_50_) and safe dose (SD) of ENR throughout the trial period, indicating that ENR had an ability of permeating the BBB of crucian carp. Evans blue (EB) dispersively distributed in brain tissue of fish exposed to SD ENR, but not in the vehicle-treated fish brain, which suggested structural failure of BBB by ENR. Expression levels of BBB-related molecules, occludin and P-glycoprotein (P-gp), were significantly down-regulated by ENR. Brain injury markers, NSE, S100B and GFAP were improved that were demonstrated by elevated contents of proteins in brain and serum. According to the data of correlation analysis, it was observed that ENR residue was closely related to S100B and GFAP, indicating that S100B and GFAP could be used as evaluation indicators for brain injury of crucian carp. All findings mentioned above suggest that ENR may cause structural change of BBB, resulting in biological brain damage in crucian carp.

## 1. INTRODUCTION

Blood-brain barrier (BBB) is considered to have a highly protective effect on CNS of organisms (Pardridge 1999). In animal brain, BBB is mainly composed of brain microvascular endothelial cells (BMECs), astrocytes (ACs), pericytes (PCs) and tight junctions (TJs) of cells or protein skeletons (Kadry et al., 2020). BBB has almost 100% barrier effect on macromolecular drugs, trans-BBB transport of macromolecules is generally through receptor-mediated endocytosis (Zhang et al., 2001). However, under normal circumstances, a small number of lipophile small molecules can directly diffuse across BBB and enter cerebrospinal fluid (CSF) without special conditions (Pardridge, 2015).

The intercellular insertion of TJs into BMECs also provides a protective effect, separating the lumen portion from the basolateral region. Claudins and Occludins are extracellular components of TJs, and key molecules of BBB. TJs is a kind of protein complex mainly formed by the interaction of a variety of transmembrane and cytoskeletal proteins, including claudin-1/5, occludins, zonula occludens-1 (ZO-1), E-cadherin and actin, and plays an important role in biological barriers (Ward et al., 2019; Wolburg et al., 2002).

Under physiological conditions, during effective screening of CSF and choroid plexus in BBB, CSF produces small amounts of proteins that are usually retained in BBB without diffusion. BBB damage can be predicted by measuring the levels of these proteins in CSF. These brain-derived proteins can be used as markers of BBB integrity (Neuwelt et al., 1999). The calcium-binding protein S100 Beta (S100B) is mainly distributed in ACs and oligodendrocytes, and had high nerve specificity. Neuron specific enolase (NSE) is expressed in neuron cytoplasm, peripheral nerves, endocrine system, platelets and red blood cells, and is mainly concentrated in neurons (Li et al., 2017). Glial fibrillary acidic protein (GFAP) highly expressed in ACs is a kind of skeletal protein with supporting function, supports adjacent neurons and BBB (Zhang, 2021). The three markers can be detected in serum if BBB is damaged (Helmrich et al., 2020). Detection of brain-derived proteins in serum is considered to be an effective method to evaluate the degree of BBB damage. Different from TJs, brain-derived proteins can not only reflect the permeability of BBB, but also reflect the damage degree of brain exposed to drug (Helmrich et al., 2020).

Enrofloxacin (ENR) is widely used in animal disease control. Its common products include raw powder, sodium salt, hydrochloride and lactate. However, the antibiotic residue and toxicity to organisms have attracted more attention, such as hepatorenal toxicity (Zhai and Li, 2017; Chen et al., 2019). In aquatic organisms, especially in fish, ENR can accumulate in fish, which impacts growth and development (Chen et al., 2019). Presently, as a member of fluoroquinolones (FQLs), ENR toxicity research mainly focuses on liver, kidney and edible animal tissues, effect on brain tissue or central nervous system (CNS) is still unknown.

Our previous work investigated the toxicity of difloxacin, another member of FQLs, to *Carassius auratus gibelio*, and found that both difloxacin doses, 2480 and 20 mg/kg, could lead to abnormal expression of inhibitory neurotransmitter γ-aminobutyric acid related anabolic genes, indicating that difloxacin influenced central nervous system (Ruan et al., 2014). ENR has most of the physical and chemical properties of FQLs, such as high lipophilicity, long biological retention, and low degradation rate (Wei et al., 2007). So it can be assumed that ENR maybe mediate impact on brain or CNS.

Pengze crucian carp (*Carassius auratus* var. Pengze), an economic species in China, was selected as the experimental object to explore the effects of ENR on the BBB permeability in fish.

## 2. MATERIALS AND METHODS

### 2.1. Experimental animals

The fish (83.60 ± 3.72 g), healthy and free from parasites, were purchased from a farm in Pengze county, Jiangxi Province, and temporarily raised for two weeks with plenty of oxygen which was pumped into the water. The feeding frequency was twice per day. During the experiment, the aquaculture water was aerated by artificial aerator, and the dissolved oxygen was stabilized at 6.50 ± 0.50 mg/L, and the water temperature and pH were controlled at 25 ± 1□ and 7.50 ± 0.30. Feeding was stopped 24 h before the experiment. Fish were divided randomly into three groups, control group (fish were orally administrated by normal saline), LD50 group (fish were orally administrated by 1949.84 mg/kg ENR) and SD group (fish were orally administrated by 194.98 mg/kg ENR), respectively (Zhao et al., 2013).

### 2.2. Determination of ENR residue by HPLC

5.0 g sample and 30 g anhydrous sodium sulfate were homogenized in 30 mL acidified acetonitrile with a high-speed tissue masher. The homogenate sample was placed in a triangular flask with glass beads (2∼3 particles/bottle), shook on a shaker for 15 min (120 rpm), centrifuged at 4500 rpm for 15 min. The supernatant was mixed with 30 mL acidified acetonitrile, centrifuged at 4500 rpm for 15 min to harvest supernatant.

25 mL n-hexane was added into the supernatant and shaken for 5 min. The lower layer of acetonitrile was transferred to the flask and evaporated to dry at 55°C. The residue was fully dissolved with 1.0 mL mobile phase and transferred to a 1.5 mL centrifuge tube, centrifuged at 4500 rpm for 5 min. The supernatant was filtered by 0.45 μm microporous membrane, and the filtrate was subjected to HPLC assay according to the Announcement of Ministry of Agriculture and Rural Affairs of China [No. 783] (2006).

### 2.3. Evans blue staining

After the OCT embedding agent infiltrates the tissue, the tissue was frozen and embedded on the quick-freezing table. Tissue was sectioned after the OCT embedding agent turned white and hardens. The sample was fixed on the slicer, the tissue surface of the sample was trimmed and then finely cut, and the thickness of the section was adjusted to 8-10 μm. After the sections were transferred to a clean slide for labeling, the red evans blue (EB) distribution (EB appears red under green light) was observed under the confocal laser scanning microscope at 405 nm excitation wavelength.

### 2.4. Concentration detection of ENR in brain tissue

The weight of crucian carp brain tissue was accurately weighed. After homogenizing the crucian carp brain tissue, 3 mL PBS and 3 mL 60% trichloroacetic acid were added. After fully mixing with a vortex oscillator, the tissue was left standing at room temperature for 30 min. After centrifugation at 3000 r/min for 10 min in a low temperature high speed centrifuge, the supernatant was harvested. A series of standard working solution of 200μL and sample supernatant were added into the 96-well plate. The absorbance at 632 nm was measured by setting the blank well to zero in the plate.

### 2.5. Quantitative polymerase chain reaction (**qPCR**)

The primers (β-actin, S100B and GFAP) for qPCR were listed in table 1, and β-actin was used as the reference gene. The reactions were performed in a thermocycler (Bio-Rad, USA) with the following profile: one cycle for 30 s at 95 □, followed by 39 cycles for 5 s at 95 L and 30 s at corresponding temperature (Table 1), and 10 s at 95 □, and then 5s at 65 L, then 5s at 95 □. The qPCR was performed in the CFX-96 real-time PCR system (Bio-Rad, USA). The qPCR reactions with 20 μL total volume per reaction (10 μL TB Green Premix, 6.4 μL double distilled water, 0.8 μL of each primer, and 2 μL cDNA templates) were set up according to the manufacturer’s protocol. Relative gene expression was analyzed using the 2^−ΔΔCt^ method.

**Table 1.**
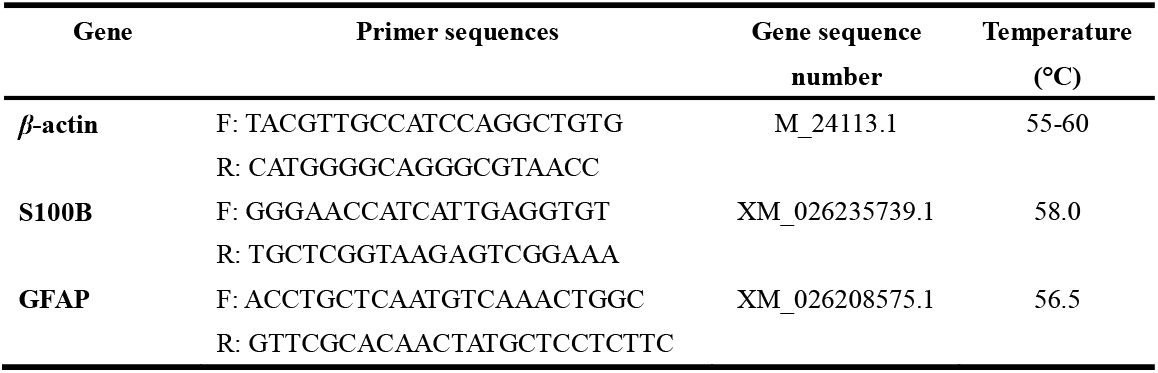
Information of qPCR primers

### 2.6. Western blot

Brain tissue homogenate containing RIPA and PMSF was centrifuged at 4□, 12000 rpm for 10 min. Supernatant was determined total protein concentration by BCA method, mixed with loading buffer, and then denatured for 10 min by boiling water bath. Samples were subjected to SDS-PAGE, and transferred to PVDF membrane. Following treatment with methanol and skimmed milk powder solution, membrane was probed with primary antibody (1:1000). After washing with PBST, membrane was incubated in labeled secondary antibody solution (1:500), and then exposed to chemiluminescence fluid in a dark room followed by analysis with Bio-Rad gel imaging system.

### 2.7. Detection of serum markers of brain injury

According to ELISA manual for experimental operation (MEIMIAN Biological Reagent Co., LTD, Wuhan), serum markers of brain injury were determined.

### 2.8. Data analysis

Data were expressed as mean ± SD, and SPSS 17.0 was used for statistical analysis. One-way ANOVA and LSD were used to make multiple comparisons for evaluating the statistical significance. *P* < 0.05 was taken as statistical significance.

## 3. RESULTS

### 3.1. ENR can pass BBB in brain crucian carp

According to the Figure 1, ENR residues in brain tissues from the two treated groups of fish were significantly different, the peak values of ENR were 712.46 ± 14.18 μg/g in LD_50_ group and 31.04 ± 0.36 μg/g in SD group, respectively. In brain tissue of LD_50_ treated fish, ENR underwent rapid biotransformation, while in brain of SD dose exposed animals, the elimination rate of ENR was slow. The data suggest that ENR can diffuse across BBB into brain tissue.

**Figure 1.**
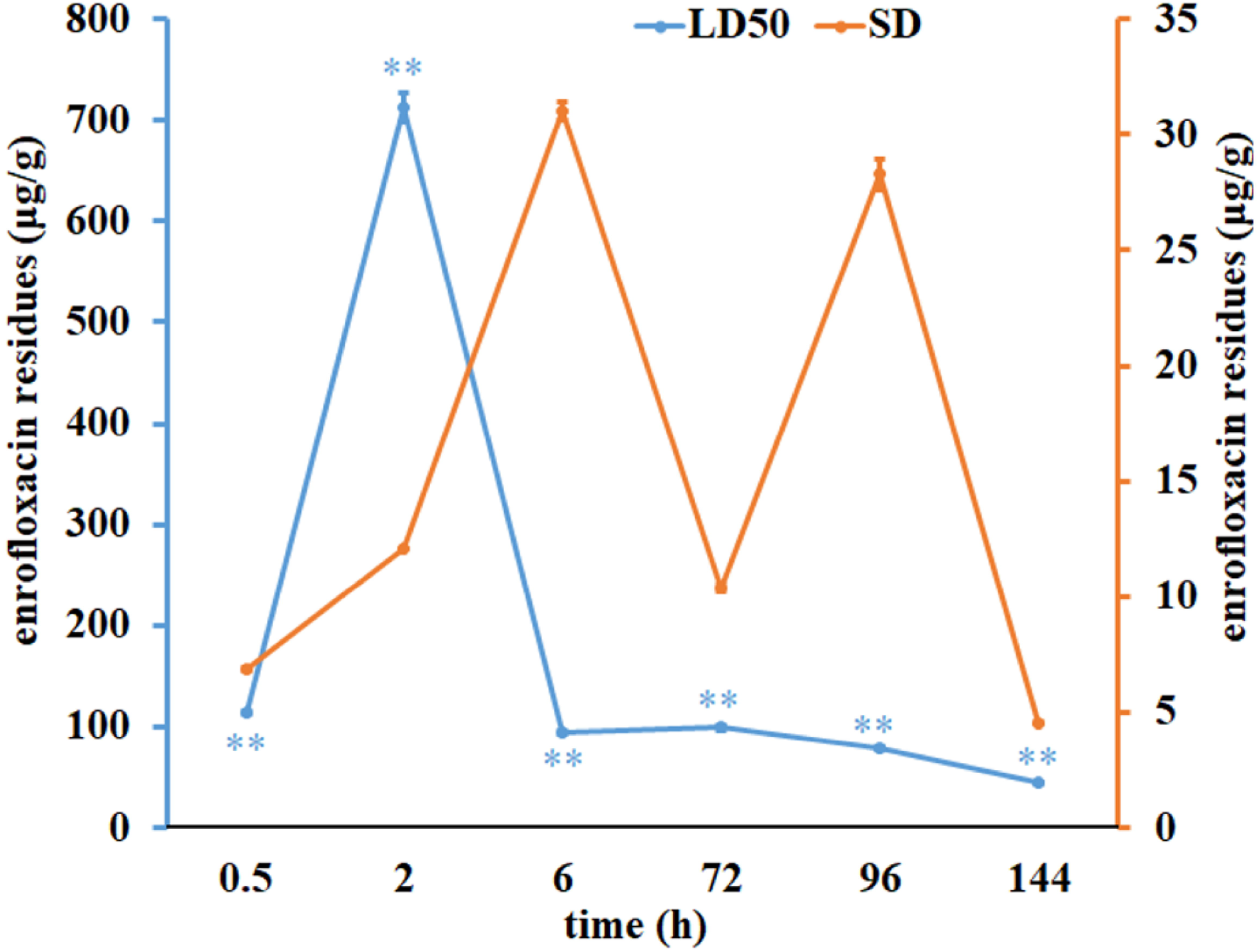
Residual amount of ENR in the brain of crucian carp (n=10). ** expresses *P*< 0.01 between LD_50_ group and SD group.

### 3.2. ENR promotes BBB permeability to EB in crucian carp

Cerebral slices of crucian carp in the control group showed that EB was clearly visible in brain microvessels (Figure 2), and the contour boundary of vascular was clear, which verified that EB did not penetrate brain microvessels into glial cells. In Figure 2, the permeability of brain microvessels was increased after ENR treatment, and EB was diffused in the brain microvessels of crucian carp, indicating that EB diffused into the glial cells. As shown in Figure 2, after treatment with ENR of SD, EB content in crucian carp brain was significantly increased (*P* < 0.01).

**Figure 2.**
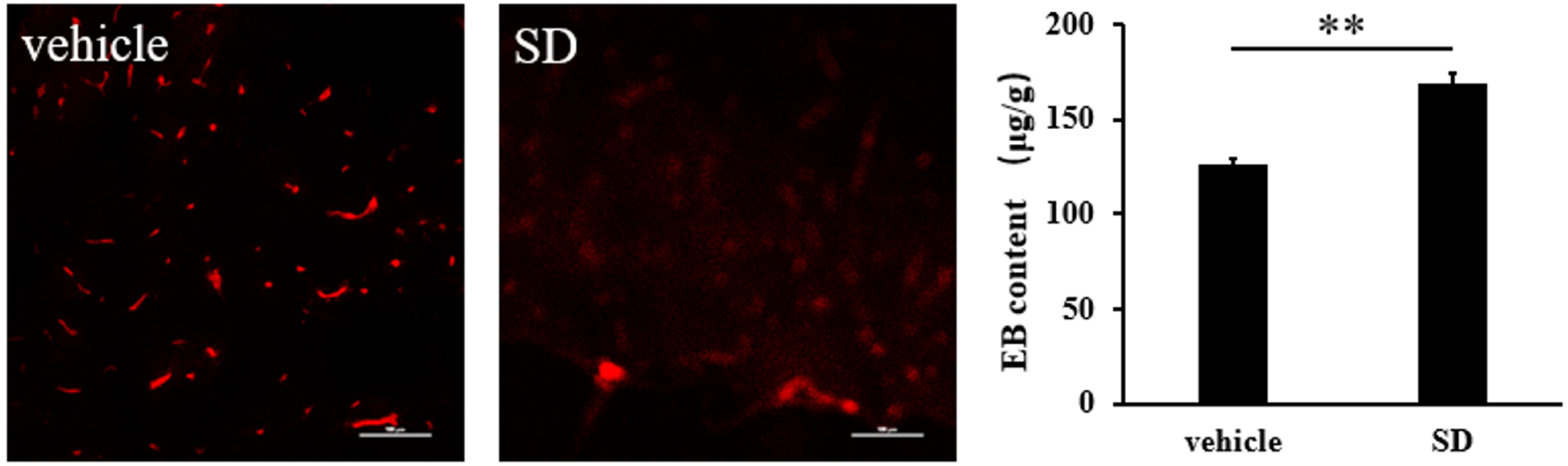
Distribution and content of EB in brain of crucian carp. EB content in the brain of crucian carp after treatment with SD of ENR for 96 h. ** expresses *P* < 0.01.

### 3.3. ENR represses expressions of occludin and P-gp, but promotes expression of GFAP, NSE and S100B in crucian carp brain

Data in Figure 3 present that contents of crucian carp BBB related proteins, occludin and P-gp, were significantly down-regulated by single oral administration of ENR at LD_50_ and SD (*P* < 0.01), moreover LD_50_ ENR was observed more obvious inhibition than SD ENR (*P* < 0.01). Meanwhile, ENR exposure elevated protein contents of GFAP and NSE in animal brain (*P* < 0.01 or *P* < 0.05), LD50 ENR was more effective in improving expression of NSE than SD ENR, significant difference between LD50 treated and SD treated fish was observed in NSE (*P* < 0.01), not in GFAP (*P* > 0.05). Apart from augment of protein content, ENR induced expression of GFAP was supported by increased transcript product (*P* < 0.01, Figure 4).

**Figure 3.**
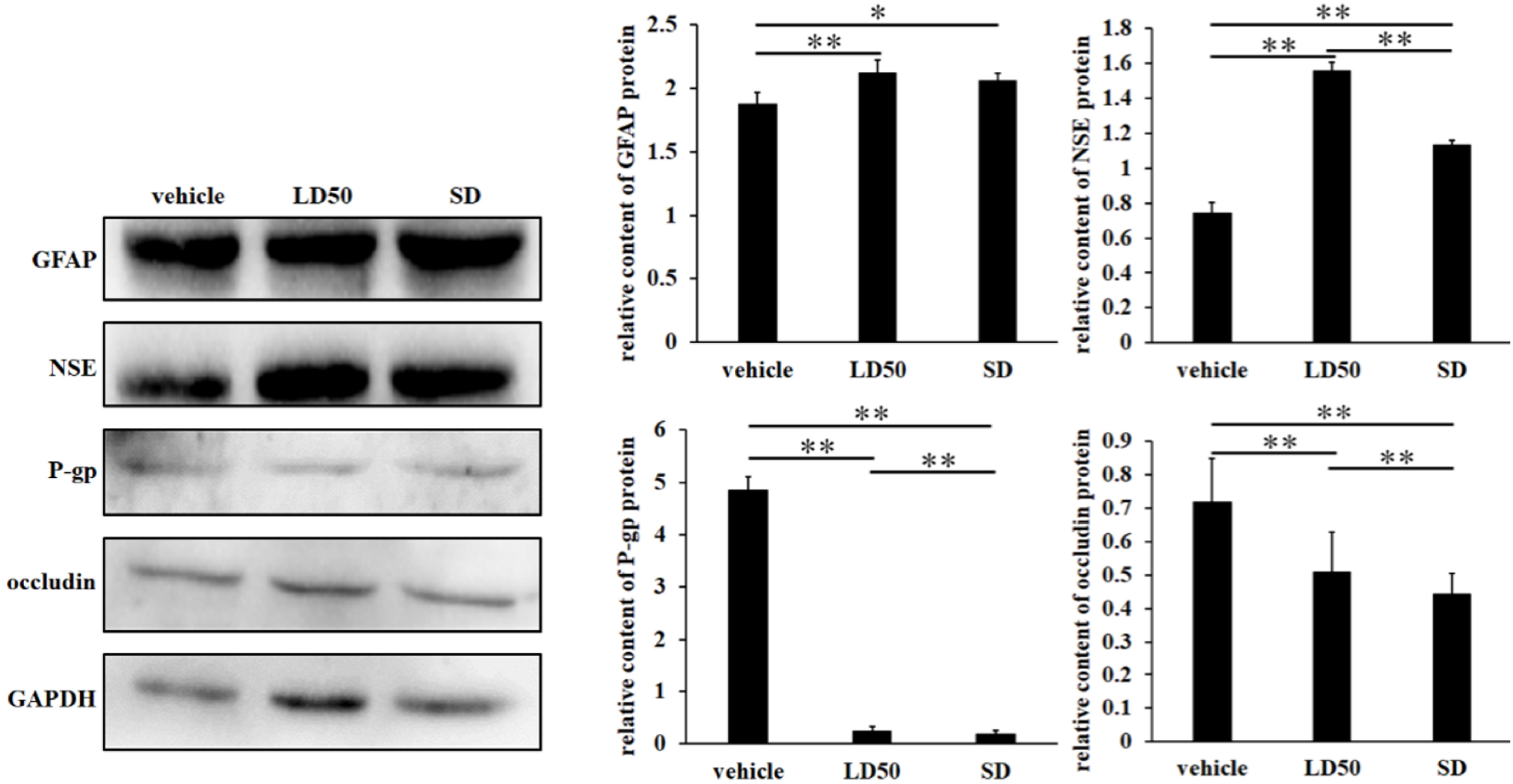
Expressions of BBB-related protein in the brain of crucian carp. Protein contents were determined by western blot analysis. * or ** expresses *P* < 0.05 or *P* < 0.01, respectively.

**Figure 4.**
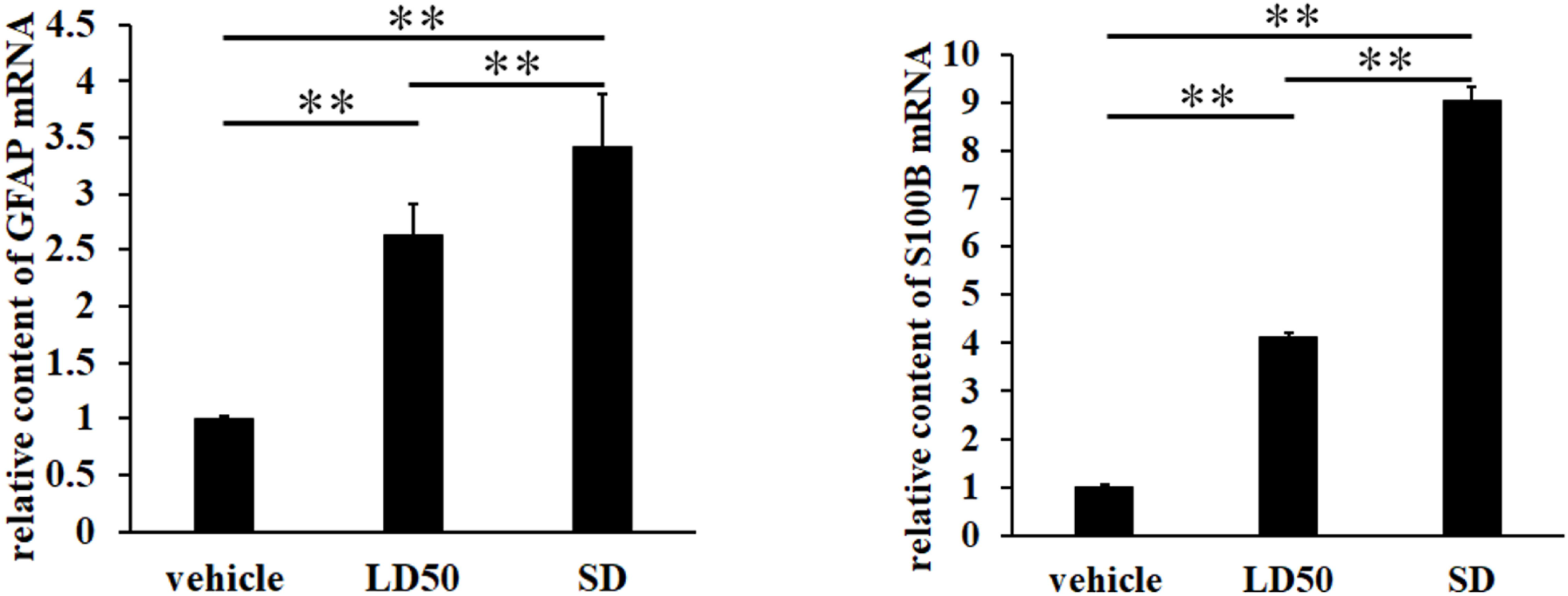
Expression of S100B and GFAP genes in crucian carp. Transcriptional levels were assessed by qPCR method. ** expresses *P* < 0.01.

Detection of transcript products also revealed that S100B was promoted by ENR (*P* < 0.01, Figure 4), the translation level could be reflected by serum content of S100B, markedly high S100B protein content was observed after treatment with ENR (*P* < 0.01, Figure 5). Data in Figure 5 also showed that ENR increased NSE and GFAP protein contents in serum. LD_50_ ENF caused extremely significant increase of GFAP protein content in serum of crucian carp (*P* < 0.01), but the change of GFAP protein content in SD group was unstable, and the difference of GFAP protein content between the two groups was extremely significant (*P* < 0.01).

**Figure 5.**
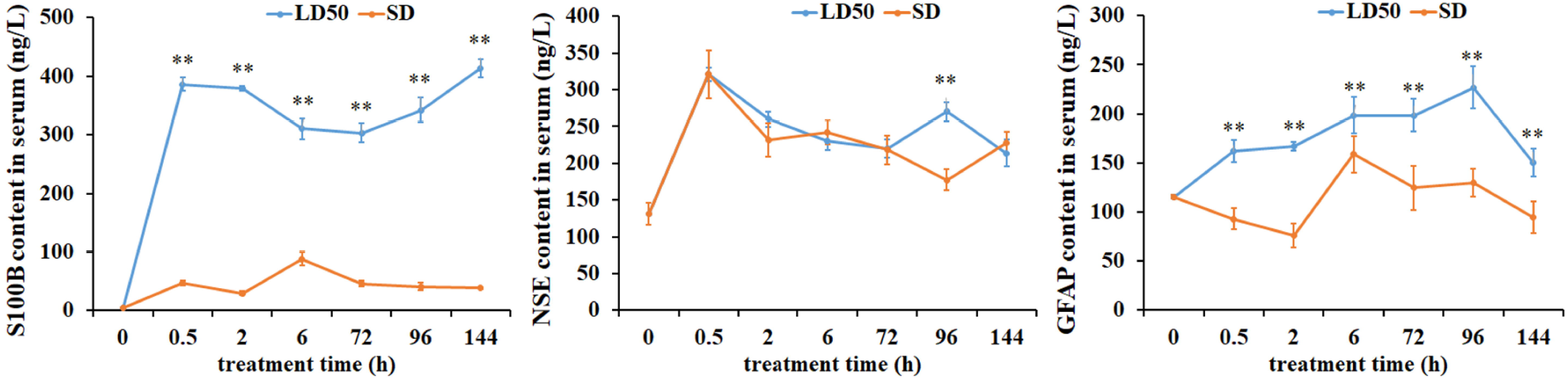
Serum contents of brain injury markers in crucian carp. Protein contents were determined by ELISA detection kits. ** expresses *P* < 0.01.

### 3.4. Correlation analysis between ENR residues and serum S100B, NSE and GFAP

Data in Table 2 present that, according to the correlation between ENR and the contents of three serum brain injury markers analyzed by Spearman rho in SPSS, the correlation between ENR and S100B, NSE and GFAP were 0.853 (*P* < 0.01), 0.140 and 0.657 (*P* < 0.05), but no close correlation was observed between ENR residue and NSE (*P >* 0.05).

**Table 2.** Correlations between ENR residues and serum S100B, NSE and GFAP

| item | correlation coefficient | significant level |
| --- | --- | --- |
| ENR residues / S100B | 0.85 | $P < 0.01$ |
| ENR residues / NSE | 0.14 | $P > 0.05$ |
| ENR residues / GFAP | 0.66 | $P < 0.05$ |

## 4. DISCUSSION

The barrier permeability of FQLs has been reported for a long time. Escudero and co-workers detected the presence of difloxacin in goat milk, demonstrating the ability of difloxacin to penetrate the hemothorax barrier into the udder (Escudero et al., 2011). It is reported that corresponding drugs can be detected in the brain tissue of rats after FQLs perfusion (Jaehde et al., 1993). In this work, ENR residue was detected in brain tissue of crucian carp exposed to ENR, two doses of LD_50_ and SD, peak values of ENR residue were 712.46 μg/g and 31.04 μg/g in LD_50_ group and SD group, respectively. As a member of FQLs, ENR could stay in the brain tissue of crucian carp in large quantities, indicating that ENR possesses high BBB permeability.

EB can locate in the injured site of BBB and has advantages of quantification (Saunders et al., 2015). Structurally, permeability of BBB is mainly controlled by BMECs, TJs and matrix membranes, it is difficult for substances to penetrate through the intact BBB into brain tissue. BBB permeability is easily affected by a variety of factors. For instance, unconventional means, such as heat stress, noise and ultrasound can increase BBB permeability of mice (Ohta et al., 2020; Sun et al., 2021). Chen et al. found that intraperitoneally injected muskone could change the ultrastructure of BBB in zebrafish (Chen et al., 2020). FQLs have high lipid solubility, so they can penetrate BBB and enter the brain easily (Ruan et al., 2014). According to the findings of this work, ENR exposure promoted EB diffusion, indicating that ENR improved permeability of BBB, then resulted in EB crossing BBB. This result further confirms that ENR can cross BBB via changing the permeability of BBB. Consequently, ENR penetrates BBB of the fish via both lipid diffusion and changing BBB permeability.

BBB permeability is regulated by a variety of factors, such as protein modification site changes (Goncalves et al., 2013), temperature (Uchida et al., 2019), heavy metals (Wang et al., 2017) and drugs (Chen et al., 2020), which can affect the structure and function of BBB. Occludin is an important molecules of tight junction widely distributed in body barrier. As an important gene in TJs, occludin is highly expressed in BBB and plays key role in the regulation of BBB function (van Leeuwen et al., 2018; Zhang et al., 2010). P-gp, a transporter of BBB, plays an important role in maintaining stability of internal environment of brain (Ding et al., 2021). In this study, it was found that expression of occludin in TJs of crucian carp BBB was extremely significantly down-regulated by ENR. This result suggested that the change of BBB permeability to ENR might be mediated by inhibiting expression of occludin in crucian carp brain.

P-gp is an important transporter of BBB. Clinically, P-gp inhibitors are used to extend retention time of drugs in brain (Nobili et al., 2012). This work revealed that ENR not only reduced the expression of TJs-related gene occludin, but also significantly inhibited P-glycoprotein, resulting in more ENR existed in brain tissue of Pengze crucian carp and consequent brain injury. S100B, NSE and GFAP have been recognized as marker molecules for astrocytes and neurons in brain, so their expression levels reflected the degree of brain injury. Our findings showed that ENR improved expression of S100B, NSE and GFAP in crucian carp brain, and higher contents of them were observed in serum. This result suggests that ACs are in a state of activation. ACs can be regulated by some drugs. Resveratrol is beneficial for brain injury caused by oxygen and sugar deprivation via inhibiting expression of S100B and GFAP, and consequent activation inhibition of ACs (Liu et al., 2020). Sevoflurane can induce expression of GFAP and activation of ACs and, resulting in neurological disorders in mice (Zhang, 2020).

## 5. CONCLUSION

In the present study, we firstly revealed that ENR improved BBB permeability of crucian carp by mediating structure disruption, consequently more ENR entered brain tissue through BBB. Elevated ENR residue induced brain injury was related to activation of ACs and neuron injury.

## CONFLICTS OF INTEREST

The authors declare no conflict of interest.

## ETHICAL APPROVAL

The entire experimental procedure was approved by the Animal Care Commission of the College of Animal Science and Technology, Jiangxi Agricultural University, China.

## AUTHOR CONTRIBUTIONS

Conceptualization, F.Z., J.R. and H.L.; formal analysis, F.Z., R.W., J.H. and H.Z.; methodology, F.Z. and R.W.; resources, L.W.; writing—original draft, F.Z.; writing—review and editing, J.R. and H.L.

## FUNDING INFORMATION

This work was supported by the National Natural Science Foundation of China (No. 31860735), the Integration and Demonstration of Rice and Fishery Green Ecological Comprehensive Planting and Breeding Technology (20192ACB60009), Key Project of Science and Technology Plan of Jiangxi Department of Education (GJJ190169), and the Key R & D Project of Jiangxi Province (20171BBF60056).

## Notes

### Competing Interest Statement

The authors have declared no competing interest.

